# Functional neural signatures of navigation impairment in early Alzheimer’s Disease

**DOI:** 10.64898/2026.06.24.734294

**Authors:** Marcia Bécu, Jonas Alexander Jarholm, Gøril Rolfseng Grøntvedt, Ivan Markel Krasovec, Per Selnes, Atle Bjørnerud, Sigrid Botne Sando, Tormod Fladby, Tora Bonnevie, Christian Doeller

## Abstract

Spatial navigation is impaired early in Alzheimer’s disease (AD), but the neural computations affected by AD pathology remain unclear. We combined virtual-reality navigation with fMRI in aged controls, preclinical AD, and early AD participants, relating navigation activity to cerebrospinal fluid biomarkers and genetic risk. Early AD was associated with dissociable alterations: amyloid-*β*-linked hippocampal hyperactivity during memory-demanding navigation, tau-linked reductions in hexadirectional modulation near the entorhinal cortex, and altered object-centered representations in posterior cingulate and entorhinal-adjacent regions. Object-centered changes related to how participants encoded and retrieved target locations. Hippocampal hyperactivity was the earliest, most robust effect and the only functional measure distinguishing preclinical individuals from controls. These findings connect AD biomarkers to distinct spatial computations in humans and reveal navigation-based functional signatures of neural dysfunction before overt dementia.

## Introduction

Spatial navigation is among the earliest cognitive domains affected in Alzheimer’s disease (AD), yet the neural computations through which AD pathology disrupts navigation remain poorly understood. AD pathophysiology is characterized by the accumulation of extracellular amyloid-*β* (A*β*) plaques and intracellular tau tangles, followed by synaptic dysfunction, neuronal loss, and cognitive decline (Wu et al., 2016; Maass et al., 2019). Notably, AD pathology affects regions that are central to spatial navigation, including the hippocampus and entorhinal cortex (EC), as well as connected posteromedial regions such as the retrosplenial cortex (RSC) and posterior cingulate cortex (PCC) (Braak & Braak, 1991; Igarashi, 2023). This anatomical overlap suggests that navigation-based paradigms may reveal early functional changes in AD before overt dementia (Lithfous et al., 2013; Vlček & Laczó, 2014; Coughlan et al., 2018).

Navigation depends on distributed neural codes that represent location, direction, landmarks, and spatial relationships (Moser et al., 2008; Bellmund et al., 2018; Epstein et al., 2017; Bicanski & Burgess, 2020). In the hippocampus, place-cell computations support cognitive maps and flexible, memory-guided navigation (O’Keefe & Nadel, 1979; Eichenbaum, 2017; Moser et al., 2008). In the EC, grid-cell representations provide a metric for space and support path integration across environments (Hafting et al., 2005; Moser et al., 2017). Posteromedial regions, including the RSC and PCC, contribute to landmark processing, self-to-object directional coding, and transformations between egocentric and allocentric reference frames (Auger & Maguire, 2013; Epstein & Vass, 2014; Alexander & Nitz, 2015; Bécu et al., 2025). Together, these systems form a distributed navigation network that is anatomically vulnerable to early AD pathology.

Task-based fMRI studies in AD and AD-risk groups have primarily used image-based memory paradigms. A recurring finding is hippocampal and network hyperactivity in preclinical AD, mild cognitive impairment (MCI), and *APOE* genetic risk states (Dickerson et al., 2005; Bookheimer et al., 2000; Berron et al., 2019). This hyperactivity appears to transition toward hypoactivity at later disease stages (Anastacio et al., 2022; Corriveau-Lecavalier et al., 2024), and elevated neuronal activity may promote amyloid and tau spread, potentially reinforcing a cycle of neurotoxicity (Cirrito et al., 2005; Palop & Mucke, 2010; Wu et al., 2016). However, whether such functional alterations extend to the neural computations that support active spatial navigation remains unknown.

Human fMRI can capture population-level spatial representations that resemble or complement those observed in animal electrophysiology. Virtual-reality navigation studies have revealed hexadirectional modulation reminiscent of grid-like coding (Doeller et al., 2010) and object-centered directional representations in medial temporal and posterior cortical regions (Bécu et al., 2025). These signals are particularly relevant to AD: tau pathology emerges early in the EC (Braak & Braak, 1991), grid-cell-like activity is impaired in transgenic mouse models of AD (Fu et al., 2017; Igarashi, 2023), and reduced hexadirectional modulation has been reported in young adult *APOEε*4 carriers and in ageing (Kunz et al., 2015; Stangl et al., 2018). Yet, no study has directly tested whether AD pathology disrupts representational codes for navigation in humans.

Here, we tested whether early AD pathology is associated with dissociable disruptions in the neural computations supporting human spatial navigation. Participants performed a virtual navigation task during fMRI, requiring them to encode and retrieve target locations within an arena containing an intramaze object and distal landmarks. We quantified three functional imaging measures: hippocampal task-dependent activity during memory demanding navigation, hexadirectional modulation reminiscent of grid-like representations, and object-centered directional representations. We then related these measures to AD-related cerebrospinal fluid biomarkers and *APOE* genetic risk.

Our findings reveal distinct functional signatures of early AD within the navigation network: amyloid-*β*-associated hippocampal hyperactivity, tau-associated reductions in hexadirectional coding near the EC, and altered object-centered representations in posterior-medial and adjacent medial temporal regions. These results link AD-related pathology to specific disruptions in spatial computations and suggest that navigation-based functional imaging can characterize subtle neural dysfunction before overt dementia.

## Results

### Identifying functional mechanisms disrupting spatial navigation in controls, preclinical and early AD participants

We used 3 Tesla fMRI to monitor the brain activity of 55 individuals, who learned and retrieved the locations of four targets within a circular arena defined by distal landmarks and a single intramaze landmark (Fig. 1A). During task trials, participants were shown the image of one of the targets in the cue phase. Subsequently, they navigated to where they think the target was and pressed a key (retrieval phase). They received feedback based on performance (feedback phase) and the object reappeared in its correct location (recollection phase).

**Figure 1.**
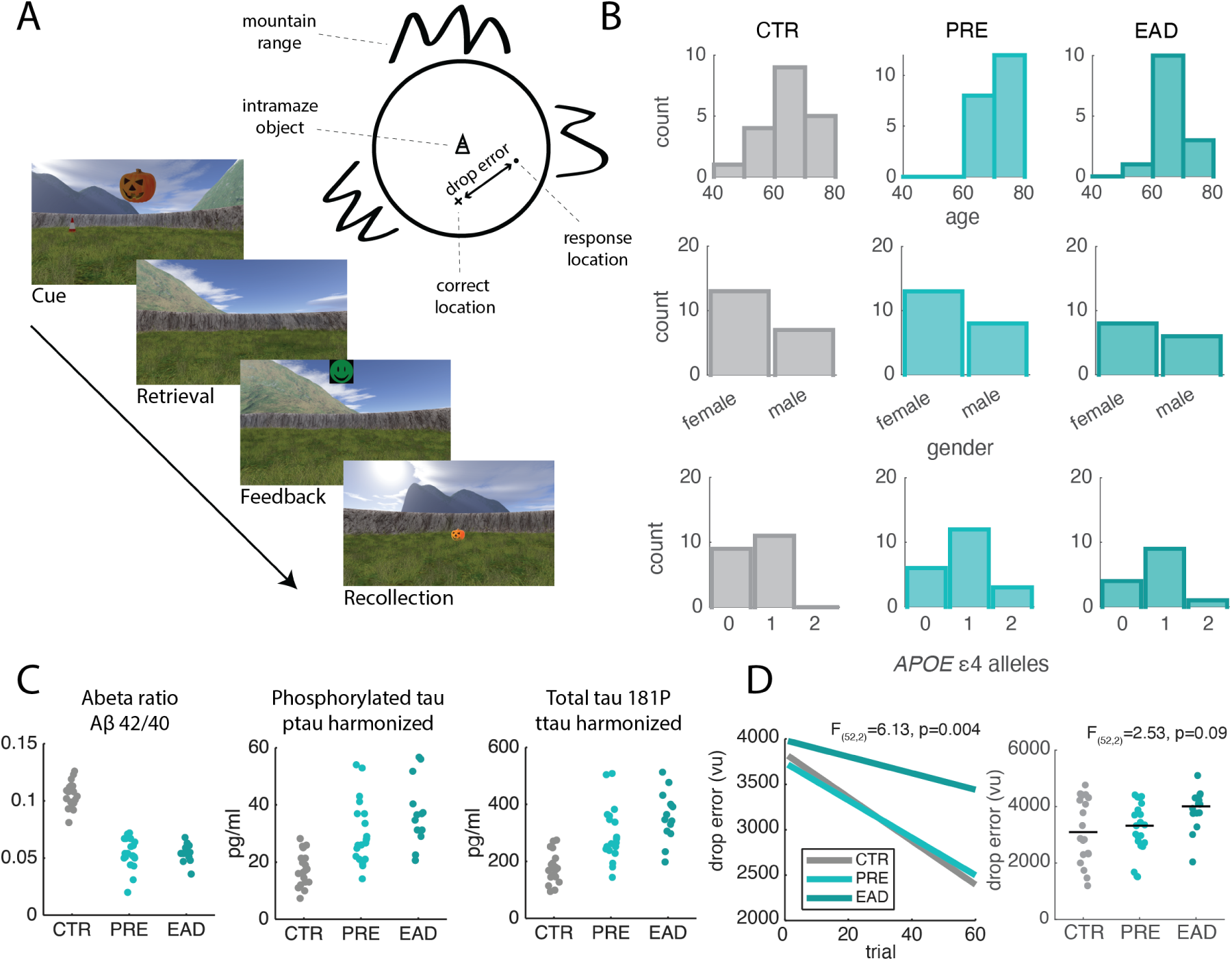
Virtual navigation task in aged controls (CTR), preclinical cases (PRE), and individuals with early Alzheimer’s disease (EAD). (A) First-person and top-down schematic view of the environment with the intramaze object (orange cone) in the center and array of distal landmarks projected at infinity in the background. (B) Histograms of age, gender and *APOEε*4 genetic status in the study groups. (C) Cerebrospinal fluid concentrations of amyloid-*β* (42/40 ratio), phosphorylated and total tau, harmonized across estimation methods (see Methods), separately for the three groups. (D) Group-wise linear fits of drop error across trials 1–60 (left) and averaged across trials (right), separately for three groups.

Participants were classified as cognitively normal or impaired by a standardized cognitive test battery (Fladby et al., 2017). Furthermore, they underwent lumbar puncture and were profiled for core cerebrospinal fluid (CSF) biomarkers of AD (A*β*42/40 ratio, phosphorylated tau, p-tau and total tau, t-tau), and *APOE* genotyped by blood samples.

The participant were further classified by a neurologist into three study subgroups: cognitively normal controls (CTR, n=20), with normal CSF amyloid biomarkers (defined by A*β*42/40 ratio) and normal cognition, preclinical AD (PRE, n=21), with pathologic CSF amyloid biomarkers and normal cognition, and early AD (EAD, n=14), with pathologic CSF amyloid biomarkers, and mild cognitive impairment.

Task performance was estimated as the drop error, defined as the distance (in virtual units, v.u.) between the response location given by the participant and the correct location (Fig. 1A). A significant group effect was observed for learning slopes (calculated using linear regression of drop error over 60 first trials; CTR slope=-24 vu/trial; PRE slope=-20.6, EAD slope=-9.1; *F*_(52,2)_ = 6.13, *p* = 0.004), with early AD participants exhibiting significantly higher slopes—indicating slower learning—compared with the two other groups (Fig. 1D, left). Although drop error generally appeared to increase with disease progression, a one-way ANOVA on drop error across study groups did not reveal a significant group difference (averaged across trials; *F*_(52,2)_ = 2.53, *p* = 0.09, Fig. 1D, right).

Following fMRI acquisition, data were analyzed using three complementary models (Supp. Fig. S1). The hyperactivity model quantified overall neural engagement during the retrieval phase, the period of highest memory demand. The hexadirectional model assessed whether activity increased when participants’ navigation aligned with data-derived six-fold axes, reflecting putative grid-like spatial coding. Finally, the object model evaluated how brain responses varied with respect to the egocentric direction of the central intramaze object. After subject-level analyses, group effects were assessed using ANOVAs designed to compare controls and participants with early AD. Whole-brain results thresholded at *p* < 0.001 (uncorrected) are reported for descriptive and exploratory purposes. Given strong a priori hypotheses regarding the involvement of the entorhinal, hippocampal, and posterior cingulate navigation network in spatial navigation and AD-related pathology, statistical inference focused on these predefined regions of interest. Multivariate analyses were subsequently used to examine associations between neural measures derived from these regions and core AD biomarkers, cognitive performance, and task-related behavioral measures.

### Hippocampal hyperactivity in Alzheimer’s disease is associated with higher amyloid-*β* concentration

To estimate memory-related fMRI activation, we modelled mnemonic phases of the task (i.e. the retrieval phase, first-level) and looked for brain areas which showed significant difference between the control (CTR) and early AD (EAD) groups (second-level). Because the human hippocampus is generally considered a task-negative area, this model explores any lack of deactivation in participants with early AD. Consistent with previous studies, we observed clusters of voxels localized in the hippocampal region, bilaterally (Fig. 2A). To further quantify this effect, we averaged the beta estimate at first-level within subject-specific masks of the hippocampi, based on Freesurfer segmentations (Fig. 2B). A two-way ANOVA (group by hemisphere) shows a significant effect of the group (*F*_(52,2)_ = 4.9, *p* = 0.01, *η*^2^ = 0.147) but no hemisphere effect (*F*_(52,2)_ = 1.13, *p* = 0.29) or interaction (*F*_(52,2)_ = 0.32, *p* = 0.72). Planned contrast comparison showed less hyperactivity in controls compared to the two other groups (CTR vs PRE: *t*_(52)_ = −2.22, *p* = 0.015; EAD vs CTR: *t*_(52)_ = 2.99, *p* = 0.002). No difference was observed between the two clinical groups (PRE vs EAD: *t*_(52)_ = −1.01, *p* = 0.84, one-tail tests, Bonferroni corrected)

**Figure 2.**
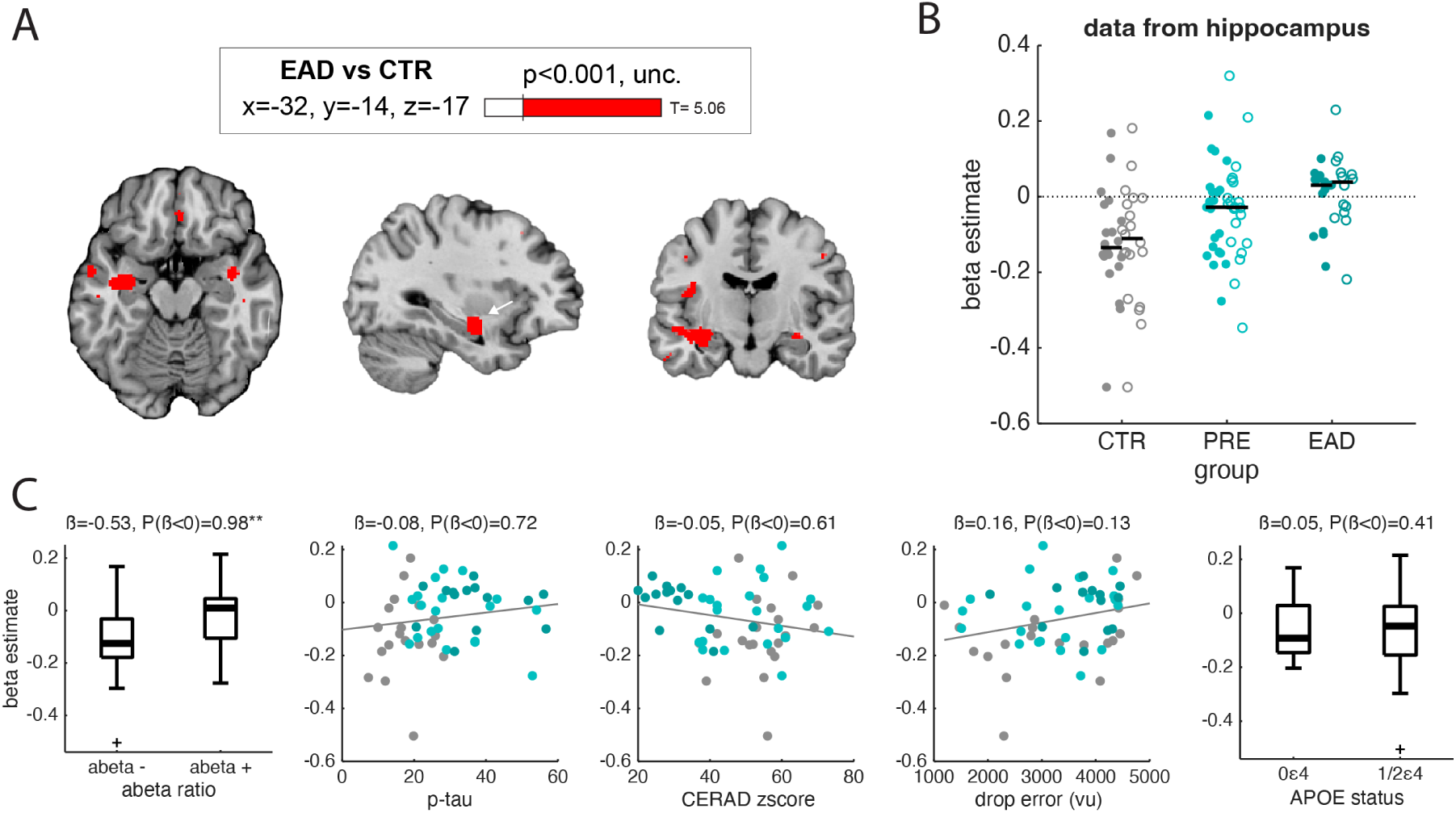
Hippocampal hyperactivity in early Alzheimer’s disease. (A) ANOVA contrast comparing controls (CTR) and early AD participants (EAD) for a whole brain parametric modulation modeling brain activity during mnemonic epochs of the virtual navigation task (retrieval phase). Clusters were identified in hippocampus and adjacent cortex, bilaterally, at uncorrected threshold (unc.). Maximum T-score, and MNI slice location. (B) Average beta estimate in the hippocampus ROI (filled: left hemisphere, empty: right hemisphere) in our three study groups. (C) Bayesian multiple regression of the hippocampal effect beta estimate (averaged across hemispheres) shows significantly more hippocampal hyperactivity in amyloid-positive participants (“abeta+”) compared to amyloid-negative participants (“abeta-”). For each regressor, we show the estimated regression coefficients (i.e. the posterior mean, *β*) and the posterior probability given the model tested (*P*(*β* < 0), negative association, *P*(*β* > 0), positive association). ** indicates strong evidence *P*(*β* > 0) or *P*(*β* < 0) > 0.975); * indicates moderate evidence (0.90–0.975); no star indicates weak or inconclusive evidence (< 0.90). For boxplots, outliers are represented by a plus sign, defined as individual values more than 1.5× the interquartile range from the lower or upper quartile. *APOE* status: 0*ε*4=non-carriers, 1/2*ε*4 = carriers of one or two *ε*4 alleles.

Bayesian multiple regression was conducted to examine associations between biomarkers, behavioral measures, and covariates with hippocampal hyperactivity (VD). The predictors included amyloid-*β* concentration (dichotomized based on our predefined disease cut-off), p-tau (entered instead of t-tau to avoid expected multicollinearity between the two variables), cognition (operationalized as the episodic memory CERAD zscore for delayed recall), task performance (drop error), and *APOE* genotype (dichotomized according to the presence versus absence of the *c*4 allele). To account for potential confounding effects, the model additionally included four covariates of no interest: volume within the ROI, functional signal quality (measured as temporal signal-to-noise ratio, tSNR, see Supp. Fig. S2), age, and sex. Directional hypotheses were evaluated using posterior probabilities (> 0.975 strong evidence; 0.90–0.975 moderate evidence; < 0.90 weak or inconclusive).

There was strong evidence for a negative association between amyloid-*β* status and hippocampal hyperactivity (β = −0.53, 95% CI [−0.96, −0.09], P(β < 0) = 0.98), indicating higher hyperactivity in amyloid-positive individuals (Fig. 2C), controlling for all other variables. Moderate evidence was observed for a positive association with male sex (β = 0.33, 95% CrI [−0.06, 0.71], P(β > 0) = 0.92), with some overlap on zero, where male participants showed increased hippocampal hyperactivity (Supp. Fig S3). All other predictors, including p-tau, *APOE* genotype, episodic memory performance (CERAD z-score), and task performance showed weak or inconclusive evidence (posterior probabilities < 0.90). Full posterior estimates are provided in Supplementary Table S1.

### Decreased hexadirectional signal in early Alzheimer’s disease correlates with phosphorylated tau concentration

Second, we aimed to estimate hexadirectional modulation in our participant sample by identifying brain voxels that exhibited increased activity when participants’ navigation trajectories aligned with empirically derived six-fold axes (Doeller et al., 2010). Straight trajectory segments during active navigation were divided into two halves (Fig. 3A, top). The first half was used to estimate the hexagonal axes orientation within the 60^◦^ space (Fig. 3A, bottom left), and the second half was used to parametrically test whether brain activity was greater along the inferred hexadirectional axes (Fig. 3A, bottom right). Comparisons between early AD patients and control participants revealed no significant differences within the medial temporal lobes (Supp. Fig. S4A). Given that a hexadirectional modulation is thought to decline with normal aging (Stangl et al., 2018) and in individuals at genetic risk for AD (Kunz et al., 2015), we next examined whether hexadirectional modulation could be detected in the medial temporal lobes of controls only.

**Figure 3.**
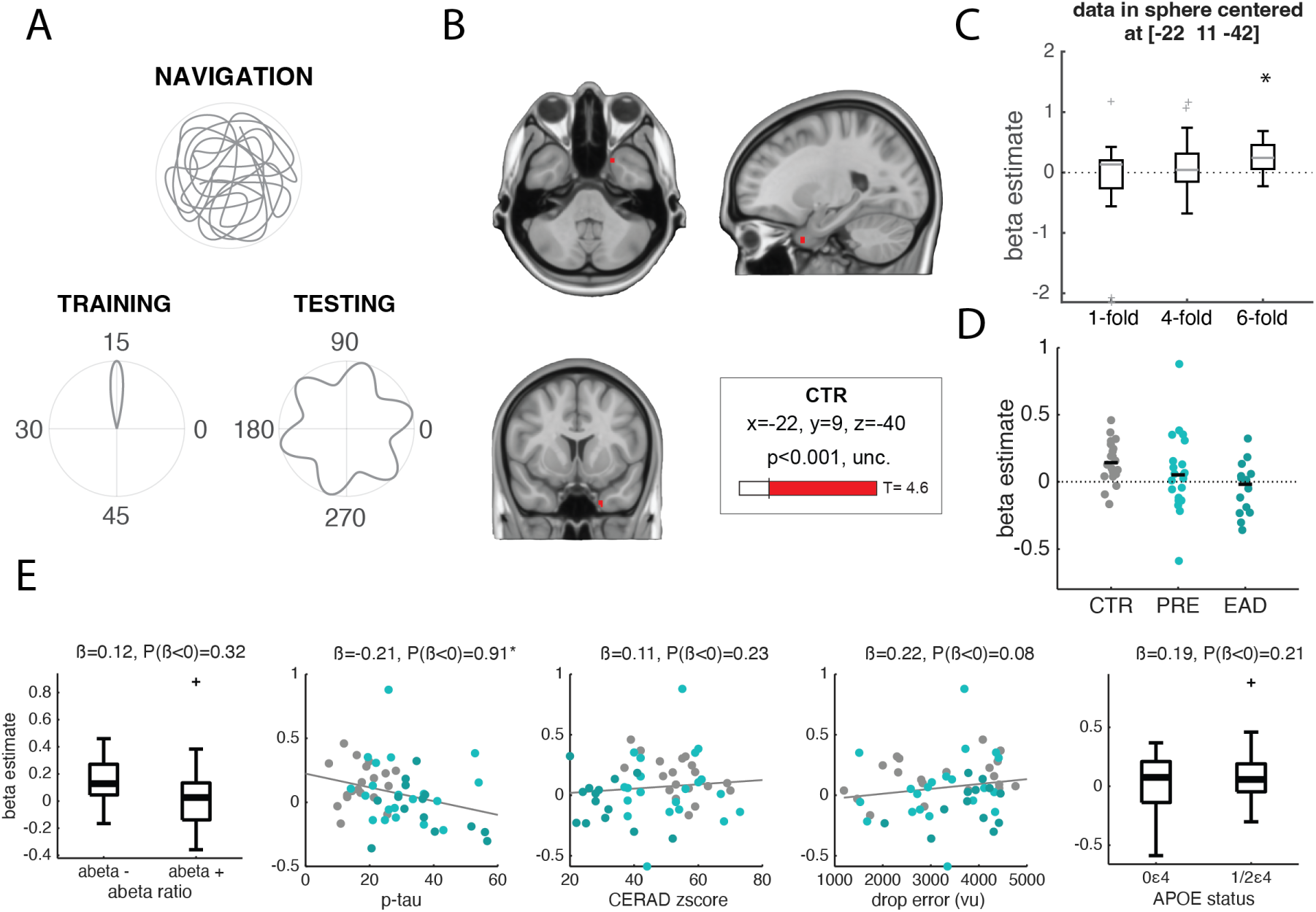
Hexadirectional activity in early Alzheimer’s disease and aged controls. (A) Model explanation: segments of active navigation were separated in two. The first half of the data was used to estimate subjective hexagonal axes orientation into the 60^◦^ space (Training), while the other half was used to parametrically test whether brain activity was higher along the hexadirectional axes (Testing). (B) Whole brain activation map of hexadirectional activity in controls only: a cluster of 14 voxels was identified close to EC, although anteriorly, in the left hemisphere, at uncorrected threshold (unc.). Maximum T-score, and MNI slice location. (C) Average beta estimate in a 1.5-voxel radius sphere centered on the control group local maxima ([−22, 11, −42]) in controls for models testing the 1*^st^*, 4*^th^* and 6*^th^* symmetries. The star indicates a significant difference from zero after Bonferroni correction for multiple comparisons. (D) Average beta value in the sphere across study groups. (E) Bayesian modeling of the hexadirectional effect shows significantly higher activity in participants with lower phosphorylated tau concentration. For each regressor, we show the estimated regression coefficients (i.e. the posterior mean, *β*) and the posterior probability given the model tested (*P*(*β* < 0), negative association, *P*(*β* > 0), positive association). ** indicates strong evidence *P*(*β* > 0) or *P*(*β* < 0) > 0.975); * indicates moderate evidence (0.90–0.975); no star indicates weak or inconclusive evidence (< 0.90). For boxplots, outliers are represented by a plus sign, defined as individual values more than 1.5× the interquartile range from the lower or upper quartile. *APOE* status: 0*ε*4=non-carriers, 1/2*ε*4 = carriers of one or two *ε*4 alleles.

In this group, we identified a significant cluster located anterior to the EC (Fig. 3B), in the left hemisphere. Although this effect was not localized directly within the entorhinal cortex (Supp. Fig. S4B), as commonly reported in the literature, we nevertheless explored it further by averaging beta estimates within a sphere centered on the peak of the control group-level cluster at MNI coordinates [−22, 11, −42].

In controls, the effect was specific to a 6-fold symmetry and was not observed for control symmetries (one-tailed Wilcoxon signed-rank tests: 6-fold, *W* = 190, *p* < 0.0003; 4-fold, *W* = 119, *p* = 0.90; 1-fold, *W* = 116, *p* = 1; Bonferroni corrected for multiple comparisons, Fig. 3C). Exploratory sphere-averaged beta estimate comparison showed significant group difference (*F*_(52,2)_ = 3.28, *p* = 0.045, *η*^2^ = 0.11), particularly between controls and early AD participants (EAD vs CTR: *t*_(52)_ = −2.562, *p* = 0.0067; CTR vs PRE: *t*_(52)_ = 1.272, *p* = 0.10; PRE vs EAD: *t*_(52)_ = 1.43, *p* = 0.078, one-tailed tests, Bonferroni corrected, Fig. 3D).

Bayesian regression analyses predicting hexadirectional modulation (VD) provided moderate evidence for an association with p-tau, such that higher p-tau was associated with lower hexadirectional modulation (β=-0.21, 95% CrI [−0.47, 0.05], P(β < 0)=0.91, Fig. 3E). Note that there was also moderate evidence that higher drop error was associated with increased hexadirectional modulation, contrary to our hypothesized effect direction (β=0.22, 95% CrI [−0.04, 0.47], P(β > 0)=0.08). For all other predictors (A*β* ratio, *APOE* status, CERAD zscore, age, sex, tSNR, and atrophy), posterior probabilities did not provide evidence for the hypothesized directional effects (all posterior probabilities ≤ 0.90, see Supp. Fig S4 and Supp. Tab. S2).

To ensure that the absence of a hexadirectional effect in early AD was not driven by subtle differences in navigation behavior that might influence how hexadirectional effects are estimated, we compared segment direction, count, and duration across the three groups. We observed no apparent differences between groups, nor any correlation with the hexadirectional effect (Supp. Fig. S5C-D, Pearson correlations between the hexadirectional effect and segment count: *r* = 0.09, *p* = 0.33, segment duration: *r* = 0.027, *p* = 0.77, segment resultant vector length: *r* = 0.003, *p* = 0.97). Furthermore, spatial stability of the hexadirectional pattern (i.e., the consistency of the estimated signal orientation across voxels within the ROI), temporal stability (i.e., the consistency of signal orientations over time), and the mean signal orientation derived from the first half of the data did not differ between groups, nor did they correlate with the hexadirectional effect (Supp. Fig. S5E-F, Pearson correlations between the effect and spatial stability: *r* = −0.052, *p* = 0.58, temporal stability: *r* = −0.058, *p* = 0.55, axes orientation: *r* = −0.03, *p* = 0.75). These results support the conclusion that the observed decline in hexadirectional representations in participants at a preclinical stage or with early AD is not attributable to differences in navigation behavior or inherent instability of hexadirectional representations, but rather reflects a genuine absence of hexadirectional modulation of brain activity.

### Disrupted object representations in posterior cingulate alter how participants encode locations within the virtual environment

We next estimated signal modulation as a function of the egocentric direction to the intramaze landmark. As expected from young adult references (Bécu et al., 2025), posterior regions should show deactivation when the object is immediately straight in front of the participant, and activity should ramp up linearly as the object moves behind.

When comparing early AD participants to controls, we found significant clusters of voxels in the posterior cingulate, bilaterally, localizing in Brodmann area 31 (Fig. 4A). Interestingly, we also observed a small cluster near the EC, close to the localization of the previously reported hexadirectional effect (MNI coordinates: [22, 10, −35], 22 voxels), though in the right hemisphere. Averaging the beta estimates for this cluster in Brodmann area 31 (Fig. 4B), a two-way ANOVA (group by hemisphere) showed a significant effect of group (*F*_(52,2)_ = 5.29, *p* = 0.008, *n^2^_g_* = 0.16) but no effect of hemisphere (*F*_(52,2)_ = 0.40, *p* = 0.53) and no interaction (*F*_(52,2)_ = 0.37, *p* = 0.69) in this data. Planned contrast comparisons, averaged across hemispheres, showed higher deactivation in controls compared to early AD (EAD vs CTR: *t*_(52)_ = 3.04, *p* = 0.002) but no difference between the other groups (CTR vs PRE: *t*_(52)_ = −0.38, *p* = 0.35; PRE vs EAD: *t*_(52)_ = −2.73, *p* = 0.99, one-tailed tests, Bonferroni corrected), suggesting weaker object representations in early AD participants.

**Figure 4.**
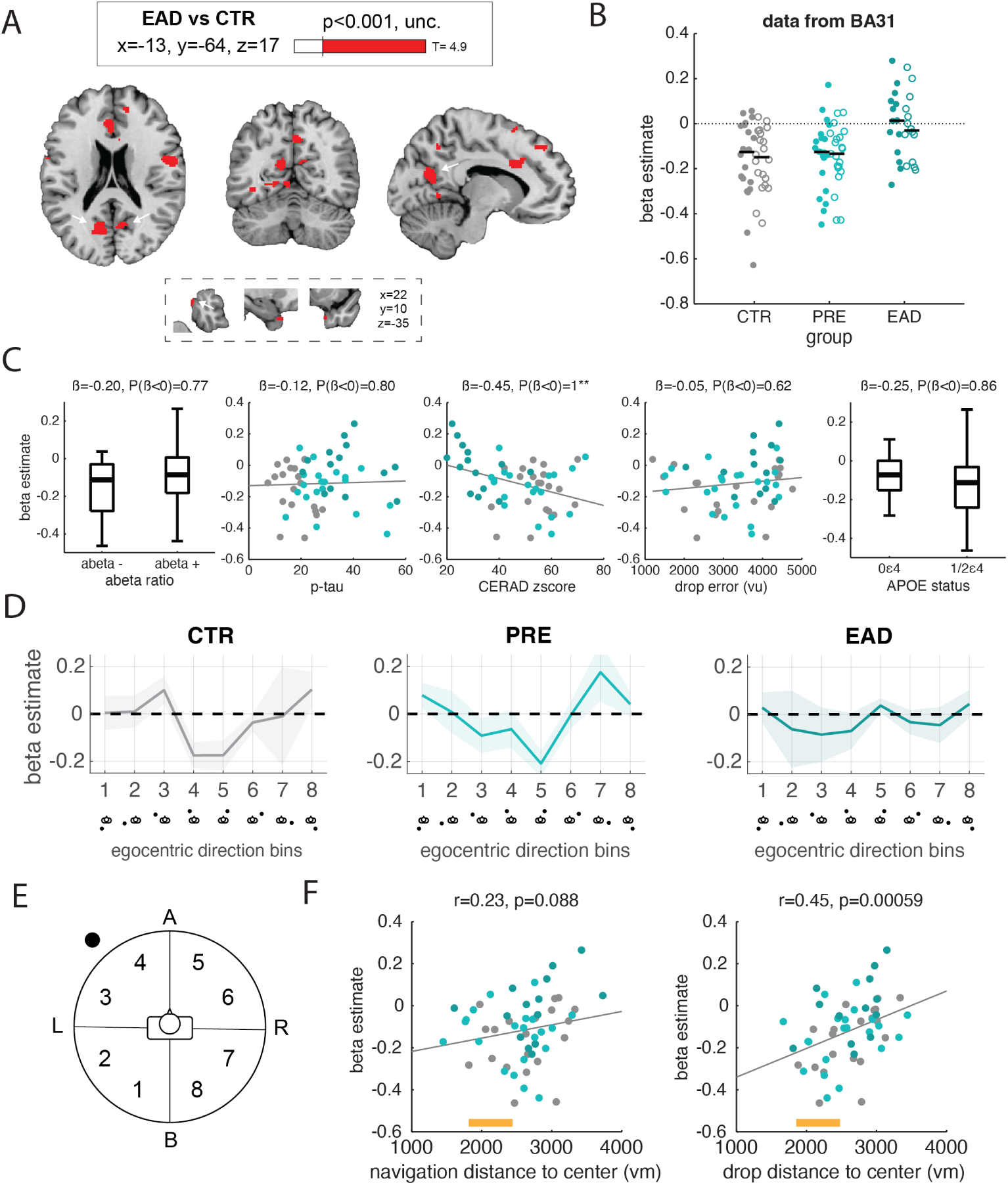
Object egocentric directional representations in early AD. (A) ANOVA contrast comparing controls (CTR) and early AD participants (EAD) for a whole brain parametric modulation modeling the egocentric direction between the participant and the central intramaze object. Clusters were identified in posterior cingulate cortex, specifically, Brodmann area 31, bilaterally, and close to the entorhinal cortex, in the left hemisphere, both at uncorrected threshold (unc.). Maximum T-score, and MNI slice locations. (B) Average beta estimate in the BA31 ROI (filled: left hemisphere, empty: right hemisphere) in our three study groups. (C) Bayesian modeling of the posterior cingulate effect (averaged across hemispheres) shows significantly lower (and therefore, better) object representations in participants with better episodic memory CERAD zscore. For each regressor, we show the estimated regression coefficients (i.e. the posterior mean, *β*) and the posterior probability given the model tested (*P*(*β* < 0), negative association, *P*(*β* > 0), positive association). ** indicates strong evidence *P*(*β* > 0) or *P*(*β* < 0) > 0.975); * indicates moderate evidence (0.90–0.975); no star indicates weak or inconclusive evidence (< 0.90). For boxplots, outliers are represented by a plus sign, defined as individual values more than 1.5× the interquartile range from the lower or upper quartile. *APOE* status: 0*ε*4=non-carriers, 1/2*ε*4 = carriers of one or two *ε*4 alleles. (D-E) Recoding every discrete directions across 8 directional regressors, we observe a scaling of the posterior cingulate response from straight ahead directions (regressors 4 and 5) to behind directions (regressors 1 and 8) in controls and preclinical participants. This relation seem to be absent in early AD cases. (F) Correlations between the object-based activity and navigation (left) and drop (right) distance to the center object location. Correct targets distance from the center object is indicated in yellow on F, as reference.

Bayesian analyses of this effect shows strong evidence for a negative association between the CERAD zscore and the outcome (β = −0.45, 95% CrI [−0.70, −0.20], P(β < 0) = 1.00, Fig. 4C) where stronger deactivation in this model was associated with better episodic memory score, controlling for all other variables. Additionally, we observed moderate evidence for a positive association with tSNR (β = 0.25, 95% CrI [0.02, 0.47], P(β > 0) = 0.96) and atrophy in the ROI (β = −0.18, 95% CrI [−0.40, 0.04], P(β < 0) = 0.92). Evidence for the remaining predictors was weak or absent, with posterior probabilities below 0.90 (see Supp. Fig S6 and Supp. Tab. S3.

To further examine how object directional representations influenced brain activity, we estimated beta weights for 8 regressors corresponding to egocentric directions relative to the intramaze object (Fig. 4D-E). While both the control and preclinical groups exhibited a deactivation for the straight-ahead direction, the early AD group showed no such modulation; in fact, the pattern was reversed, with slightly elevated activation for the straight ahead direction. We then asked whether this deficit in object direction representation also affected behavior, specifically how participants encoded target locations relative to the center object. To investigate this, we measured the distance from the center at which participants navigated and dropped the targets. Participants with higher beta estimates (indicating poorer directional object representations) tended to navigate (*r* = 0.23, *p* = 0.088, Fig. 4F, left) and drop targets farther from the center object (*r* = 0.45, *p* = 0.00059, Fig. 4F, right). Conversely, participants with better object representations navigated and dropped targets at distances consistent with the expected location relative to the center object.

Finally, we investigated whether the three functional signatures identified here were correlated with each other. Across all participants, we found that beta values for the object model in the posterior cingulate was correlated with the hexadirectional model in EC (*r* = −0.34, *p* = 0.009), but not with the hippocampal hyperactivity model (*r* = 0.13, *p* = 0.33). Similarly, the hexadirectional model and the hyperactivity model showed no significant association (*r* = −0.1, *p* = 0.47).

## Discussion

By recording brain activity while participants with early AD, biomarker-positive individuals in the preclinical stage of AD, and older control participants navigated a virtual arena, we identified three distinct pathways through which AD neuropathology disrupts the neural systems underlying spatial navigation: *(i)* amyloid-*β*–associated network hyperactivity, *(ii)* tau-related degradation of hexadirectional representations, and *(iii)* impaired object-centered representations, which compromise the use of salient environmental landmarks for anchoring spatial memory.

Hippocampal hyperactivity emerged as the most prominent and the earliest functional alteration in our sample. In controls, mnemonic task moments elicited robust deactivation of the hippocampus, whereas participants with preclinical or early AD showed reduced or absent deactivation, consistent with pathological hyperactivity. This aberrant activation pattern distinguished preclinical individuals from both cognitively normal controls and participants with MCI, highlighting hippocampal hyperactivity as a potential early functional indicator for AD. Notably, the degree of hyperactivity positively correlated with amyloid-*β* concentration measured via CSF. This relationship echoes rodent findings in which hyperactive neurons are found exclusively near amyloid plaques (Busche et al., 2008). Such coordinated hyperactivity may be associated to elevated risk of, or presence of subclinical, seizure-like activity, a phenomenon also reported in humans (Lozsadi & Larner, 2006). Our finding of hippocampal hyperactivity in individuals with preclinical and early AD is consistent with previous studies (Dickerson et al., 2005; Bookheimer et al., 2000; Berron et al., 2019), and extends this evidence to active 3D spatial navigation. We also found greater hyperactivity in male participants, in contrast to an earlier study that reported no sex-related differences in hippocampal hyperactivity (Corona-Long et al., 2020). The basis for this discrepancy remains unclear and should be examined in future work.

We identified a cluster exhibiting hexadirectional (6-fold) modulation near the entorhinal cortex (EC) in the control group. Although modest in size, this cluster displayed key properties of hexadirectional coding. The limited spatial extent of the effect is not unexpected, given that hexadirectional signals in human fMRI are often tenuous, arise in a region such as the EC that is particularly susceptible to susceptibility-related signal loss and air-tissue artefacts, and are not always robustly detected even in young adults (Kransberg et al., 2026). Moreover, EC function and hexadirectional coding are known to be affected by normal ageing and *APOE* status (Kunz et al., 2015; Stangl et al., 2018), factors that are especially relevant in the present cohort of older adults and MCI participants. The strength of this effect diminished with disease progression, aligning with classical observations that tau pathology initially accumulates in EC (Braak & Braak, 1991). Indeed, hexadirectional signal magnitude correlated with CSF tau concentration. Importantly, we verified that the absence of hexadirectional modulation in preclinical and early AD participants was not attributable to differences in navigation behavior, nor to instability of signal orientation across time or space. These controls allow us to suggest a genuine loss of hexadirectional coding in early AD. However, given the exploratory nature of the sphere-based analysis, the modest cluster size, and the relatively small effect sizes observed at both the group and tau-related levels, the findings should be considered preliminary pending independent replication. Interestingly, the hexadirectional signal we observed was located slightly anterior to EC. Prior work has also reported grid-like activity outside the EC proper (Doeller et al., 2010; Constantinescu et al., 2016). Nonetheless, we cannot exclude potential influences of distortion correction or atlas-based alignment error in this older cohort.

Finally, object-centered representations—operationalized via egocentric directional modulation—declined with disease progression, with significant reductions emerging only after the onset of objective cognitive impairment. Declining object-centered coding correlated with episodic memory performance, mirroring prior findings implicating the posterior cingulate cortex (PCC) in episodic encoding and retrieval (Papma et al., 2017; Natu et al., 2019). We thus interpret this association as reflecting shared neural substrates, rather than contamination of the egocentric model by episodic-memory demands. Although object-centered coding correlated with atrophy and fMRI signal quality within the ROI, these variables were included as covariates in the model, indicating that the relationship with episodic memory persists beyond structural and signal-level differences. Qualitatively, PCC activity decreased when the object was directly visible and increased as the object moved out of view—consistent with previously described mechanisms supporting vision-independent maintenance of salient object representations (Bécu et al., 2025). This modulation was absent in participants with early AD. While object-centered representations did not correlate with overall drop-error magnitude, they nonetheless influenced behavior. Poorer object-centered coding was associated with systematic biases in recalled target locations—specifically, repulsion from the object or attraction toward the boundary—whereas stronger object representations facilitated more accurate encoding of target distances from the central object and/or the boundary.

Kunz et al. (2015) previously proposed that impaired hexadirectional coding in young individuals at risk for AD may be compensated by heightened hippocampal activity. In our older cohort spanning normal aging to early AD, we did not observe a relationship between hippocampal hyperactivity and hexadirectional impairment. The strong association between amyloid-*β* and hyperactivity may indicate a pathological, rather than compensatory, interpretation of hippocampal hyperactivity, although confirmation will require complementary molecular, electrophysiological, or pharmacological approaches. On the other hand, hexadirectional and object-centered representations may partially rely on shared computational pathways, as both effects localized near the EC (albeit in opposite hemispheres), and these two functional markers were themselves correlated.

In sum, while all three navigation-related functional indicators—hippocampal task-dependent activity, hexadirectional representations, and object-centered representations—were associated with AD progression, hippocampal hyperactivity emerged as the earliest and most robust effect. Crucially, it was the only functional imaging measure capable of distinguishing preclinical individuals from cognitively normal controls, underscoring its promise as a sensitive functional signature for early AD across tasks. More broadly, these findings link AD-related neuropathology to dissociable disruptions in the neural computations supporting spatial navigation, offering a mechanistic bridge between systems neuroscience and the functional characterization of AD pathophysiology. Clinically, they suggest that navigation-based functional imaging indicators may help identify subtle neural dysfunction before overt cognitive impairment, with potential value for tracking disease progression and monitoring intervention effects.

## Methods

### Participants

A total of 73 participants were recruited from Akershus University Hospital, Loeren-skog and St. Olavs Hospital in Trondheim, Norway, between May 2018 and September 2020, as part of the Dementia Disease Initiation study. Among this original sample, 18 participants were excluded because of missing data (brain, behavior or biomarkers), leading to a final sample of 55 participants (range: 49-79 years, average: 68.0, median: 67.0), 34 females, 21 males). Demographic data is provided in table 1. The recruitment of the cases (n=35) was self-referred (n=25 or 71%)(media advertisements, public lectures, or referral through current participants) or referred by local outpatient or memory clinics (n=10, 29%). Controls (n=20) were self-referred (n=17, 85%) or recruited from memory clinics (n=3, 15%). All screening and experimental procedures were performed in accordance with the principles of the Declaration of Helsinki and were approved by the Norwegian Regional Committee for Medical and Health Research Ethics South East (REK South East, 01/11/2012, D2013/150). All participants were voluntary and gave their informed consent approval. Inclusion criteria for the DDI study were *i)* age 40-80 years and *ii)* Scandinavian mother tongue. Exclusion criteria were *i)* dementia, *ii)* other brain diseases of non-neurodegenerative etiology (like stroke, significant psychiatric comorbidity, traumatic brain injury, infections or inflammatory disorders of the brain).

**Table 1.**
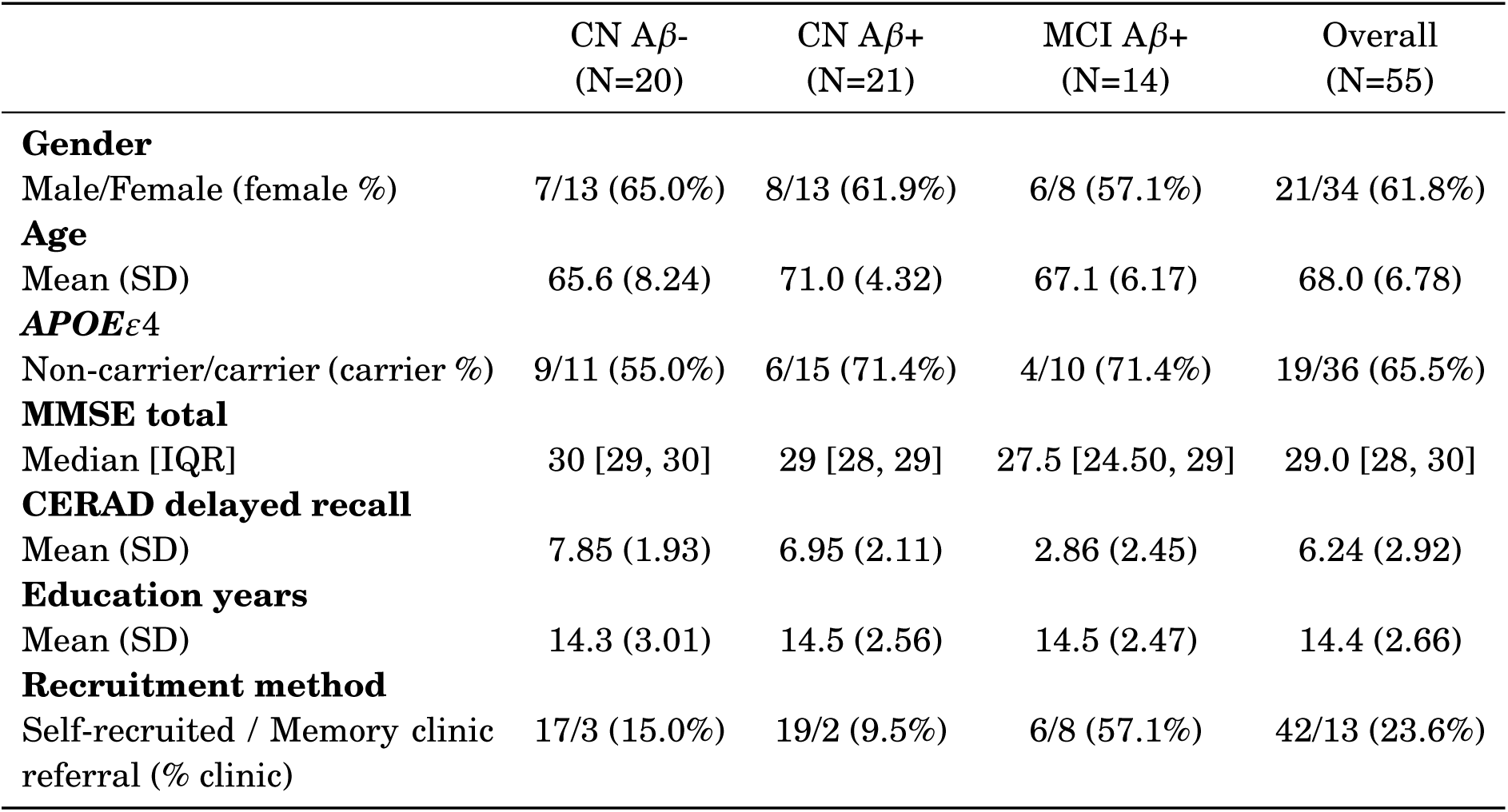
Participant characteristics. CN=cognitively normal, MCI=mild cognitive impairment, SD=standard deviation, IQR=inter-quartile range, Aβ -/+=amyloid-β negative/positive.

| | CN Aβ-<br>(N=20) | CN Aβ+<br>(N=21) | MCI Aβ+<br>(N=14) | Overall<br>(N=55) |
| --- | --- | --- | --- | --- |
| **Gender** |  |  |  |  |
| Male/Female (female %) | 7/13 (65.0%) | 8/13 (61.9%) | 6/8 (57.1%) | 21/34 (61.8%) |
| **Age** |  |  |  |  |
| Mean (SD) | 65.6 (8.24) | 71.0 (4.32) | 67.1 (6.17) | 68.0 (6.78) |
| **APOEε4** |  |  |  |  |
| Non-carrier/carrier (carrier %) | 9/11 (55.0%) | 6/15 (71.4%) | 4/10 (71.4%) | 19/36 (65.5%) |
| **MMSE total** |  |  |  |  |
| Median [IQR] | 30 [29, 30] | 29 [28, 29] | 27.5 [24.50, 29] | 29.0 [28, 30] |
| **CERAD delayed recall** |  |  |  |  |
| Mean (SD) | 7.85 (1.93) | 6.95 (2.11) | 2.86 (2.45) | 6.24 (2.92) |
| **Education years** |  |  |  |  |
| Mean (SD) | 14.3 (3.01) | 14.5 (2.56) | 14.5 (2.47) | 14.4 (2.66) |
| **Recruitment method** |  |  |  |  |
| Self-recruited / Memory clinic referral (% clinic) | 17/3 (15.0%) | 19/2 (9.5%) | 6/8 (57.1%) | 42/13 (23.6%) |

### Cognitive staging

The study clinical protocol included a complete medical history, a standardized cognitive test battery covering different cognitive domains, as recommended by the National Institute on Aging-Alzheimer’s Association (NIA-AA) working group (Albert et al., 2011) and clinical examinations including complete neurological examination and lumbar punction, as described hereafter. The examinations were performed within a time window of 3 months. However, for 14 participants, the interval between scanning and final diagnosis exceeded this time window. Nevertheless, earlier or later follow-up assessments confirmed that these participants remained cognitively unchanged at the time of scanning. We therefore considered it unlikely that their AD biomarker status had changed within this interval.

Core cerebrospinal fluid (CSF) biomarkers (A*β*_42/40_, total tau, and phosphorylated tau) were acquired by a lumbar puncture and analyzed using Enzyme-linked Immunosorbent assay (ELISA) methods. The samples were collected in sterile polypropylene tubes (Thermo Nunc) before noon. The analyses were performed at the Department of Interdisciplinary Laboratory Medicine and Medical Biochemistry at Akershus University Hospital. ELISAs based on monoclonal antibodies from Innotest (Fujirebio, Ghent, Belgium) were used for samples collected before October 2020, to determine concentration of CSF total tau (t-tau, using hTau Ag kits) and phosphorylated tau (p-tau, using 181P kits). Due to an update in laboratory equipment, the concentrations of t-tau and p-tau of samples collected after October 2020 were determined with Elecsys t-tau and p-tau kits. Tau values from the two methods were harmonized based on coefficients presented in Lifke et al. (2019). A*β*_1−42_ and A*β*_1−40_ were analyzed using the QuickPlex SQ 120 system from Meso Scale Discovery (MSD, MD, USA; V-Plex Ab Peptide Panel 1 6E10 kit, K15200E-1). These analyses were performed according to the manufacturer’s standards. See Fladby et al. (2017) for a detailed description of the estimation of biomarkers.

The cognitive assessment included the following tests: Mini Mental State Examination (MMSE) (Folstein et al., 1975), Geriatric Depression Scale (GDS) (Yesavage et al., 1982), clock drawing test (Shulman, 2000), CERAD word list (Wagle et al., 2023), Visual Object and Space Perception Battery (VOSP) silhouette identification (Warrington & James, 1991), Trail Making Test A-B (TMT) (Bowie & Harvey, 2006) and the COWAT word fluency test (Benton & Hamsher, 1989). Normative data adjusted for age and education in culturally relevant populations were used to calculate the T-scores (Kirsebom et al., 2019; Lorentzen et al., 2023; Warrington & James, 1991).

*APOE* genotyping was performed via EDTA blood samples, and analyzed at Akershus University Hospital (Gene technology division, Department of interdisciplinary Laboratory Medicine and Medical biochemistry), using real-time PCR combined with a TaqMan assay (Applied biosystems, Thermo Fisher Scientific, Waltham, USA).

Subjects were classified as cognitively normal (CN) if the performance of all cognitive tests was normal, and as mild cognitive impairment (MCI) if one or more tests were abnormal, and if the clinical criteria of dementia were not met (McKhann et al., 2011). Abnormal performance on cognitive tests was defined as ≤ 1.5 SD below the normative mean (t-score of 35 or less), as described in the DDI research protocol (Fladby et al., 2017). Subjects with amyloid pathology (A+) were defined as preclinical AD (CN A+) or prodromal AD (MCI A+). We used the ratio A*β*_42/40_ to determine cerebral amyloid pathology, based on an amyloid-PET verified cut-off presented in Siafarikas et al. (2021). Participants were classified as amyloid positive (A+) when CSF ratio A*β*_42/40_ ≤ 0.077 and amyloid negative (A-) when CSF ratio A*β*_42/40_ ≥ 0.077. For a subset of cases for which lumbar puncture could not be performed due to medical conditions or unwilling participant (n=3), we used results from amyloid-PET^18^F-flutemetamol examinations, evaluated qualitatively according to the manufacturers specifications. For some of the cases missing both lumbar puncture or amyloid-PET at the staging timepoint, we reclassified them as A+ if a former assessment (CSF puncture or PET scan) confirmed amyloid positivity, or A-if a status as amyloid negative was confirmed at a later assessment.

### fMRI acquisition

High-resolution T1-weighted images are acquired on 3T Siemens Skyra and Prisma scanners in Trondheim and Oslo, respectively (Siemens Healthcare, Erlangen, Germany) and 32-channel head coil. Anatomical scans used a MPRAGE sequence (Magnetization Prepared - RApid Gradient Echo) with following parameters: echo time=3.16 ms, repetition time=1.9 s, voxel size=1mm^3^, flip angle=9^◦^, inversion time=900 ms, slice thickness=1 mm, for a total of 192 sagittal slices. BOLD T2*-weighted functional images are acquired with single-echo EPI sequence with the following parameters: TR=1.02 s, TE=34.6 ms, flip angle=55^◦^, voxel size=2 mm isotropic, field of view (FoV)=210 mm in each direction, 66 slices, multi band acceleration factor=6. Subsequent blocks of 575 volumes are acquired. In addition, 10 volumes were acquired in opposite phase encoding direction, to correct for distortions. The time in scanner is approximately 1 hour.

### Experimental task

#### Virtual environment

The virtual environment consists of a grassy plane surrounded by a circular cliff (diameter: 9500 virtual units (vu), developed with UnrealEngine2 Runtime software, Epic Games). The background shows distal cues (mountains, clouds, sun) projected at infinity. A traffic cone at the center of the arena is used as an intramaze landmark. The first-person perspective is experienced at 128 vu above ground. Forward movement (800 vu/s) and left/right rotation are only allowed. Virtual heading and location are recorded at 10 Hz.

#### Task

During initial learning, participants collected four targets in the arena by running over them and learning their positions. The targets and their locations were randomly selected from 200 possible configurations and order for each participant. During each subsequent trial, participants were shown a picture of one targets at the top of the screen (cue phase). They were instructed to replace the targets by moving to the location they believed to be correct and to press a button to indicate their response (retrieval phase). Feedback was then provided in the form of smiley faces based on the drop error (feedback phase; very good: <1000 vu; good: 1000–2000 vu; intermediate: 2000–3000 vu; fair: 3000–4500 vu; poor: >4500 vu). Afterward, the targets appeared in its correct position and had to be recollected (recollection phase). A fixation cross was presented during a variable intertrial interval (ITI phase; 3–5 s, mean = 4 s) before the start of the next trial. The session ended automatically after 43 min; therefore, the total number of trials varied across participants. Finally, four additional trials (one per object) were conducted without feedback. This task was adapted from Doeller and colleagues (Doeller & Burgess, 2008; Doeller et al., 2010).

Before going in the scanner, participants get a training session in a different virtual environment (same boundary, different distal cues, no intramaze landmark), using the same procedure. This training session involves two targets only.

## Data preprocessing

Results included in this manuscript come from preprocessing performed using FM-RIPREP (Esteban et al., 2019) 20.2.2, which is based on NIPYPE (K. Gorgolewski et al., 2011; K. J. Gorgolewski et al., 2018) 1.6.1 (RRID:SCR_002502).

### Anatomical data preprocessing

The T1-weighted (T1w) image was corrected for intensity non-uniformity (INU) with N4BiasFieldCorrection (Tustison et al., 2010), distributed with ANTS (Avants et al., 2008) 2.3.3 (RRID:SCR_004757), and used as T1w-reference throughout the workflow. The T1w-reference was then skull-stripped with a Nipype implementation of the antsBrainExtraction.sh workflow (from ANTs), using OASIS30ANTs as target template. Brain tissue segmentation of cerebrospinal fluid (CSF), white-matter (WM) and gray-matter (GM) was performed on the brain-extracted T1w using fast (Zhang et al., 2001) (FSL 5.0.9, RRID:SCR_002823). Brain surfaces were reconstructed using recon-all (FreeSurfer 6.0.1, RRID:SCR_001847), and the brain mask estimated previously was refined with a custom variation of the method to reconcile ANTs-derived and FreeSurfer-derived segmentations of the cortical gray-matter of Mindboggle (Klein et al., 2017) (RRID:SCR_002438). Volume-based spatial normalization to one standard space (MNI152NLin2009cAsym) was performed through nonlinear registration with antsRegistration (ANTs 2.3.3), using brain-extracted versions of both T1w reference and the T1w template. The following template was selected for spatial normalization: ICBM 152 Nonlinear Asymmetrical template version 2009c (Fonov et al., 2009) (RRID:SCR_008796; TemplateFlow ID: MNI152NLin2009cAsym).

### Functional data preprocessing

For each BOLD runs, the following preprocessing is performed. First, a reference volume and its skull-stripped version were generated using a custom methodology of fMRIPrep. BOLD runs were slice-time corrected using 3dT-shift from AFNI (Cox & Hyde, 1997) 20160207 (RRID:SCR_005927). Head-motion parameters with respect to the BOLD reference (transformation matrices, and six corresponding rotation and translation parameters) are estimated before any spatiotemporal filtering using mcflirt (Jenkinson et al., 2002) (FSL 5.0.9). A B0-nonuniformity map (or fieldmap) was estimated based on two (or more) echo-planar imaging (EPI) references with opposing phase-encoding directions, with 3dQwarp (Cox & Hyde, 1997) (AFNI 20160207). Based on the estimated susceptibility distortion, a corrected EPI (echo-planar imaging) reference was calculated for a more accurate co-registration with the anatomical reference. The BOLD reference was then co-registered to the T1w reference using bbregister (FreeSurfer) which implements boundary-based registration (Greve & Fischl, 2009). Co-registration was configured with six degrees of freedom. The BOLD time-series (including slice-timing correction when applied) were resampled onto their original, native space by applying a single, composite transform to correct for head-motion and susceptibility distortions. These resampled BOLD time-series will be referred to as preprocessed BOLD in original space, or just preprocessed BOLD. Several confounding time-series were calculated based on the preprocessed BOLD: framewise displacement (FD), DVARS and three region-wise global signals. FD was computed using two formulations following Power et al. (2014) (absolute sum of relative motions) and Jenkinson et al. (2002) (relative root mean square displacement between affines). FD and DVARS are calculated for each functional run, both using their implementations in Nipype (following the definitions by Power et al. 2014). The three global signals are extracted within the CSF, the WM, and the whole-brain masks. Additionally, a set of physiological regressors were extracted to allow for component-based noise correction (Behzadi et al., 2007) (CompCor). Principal components are estimated after high-pass filtering the preprocessed BOLD time-series (using a discrete cosine filter with 128 s cut-off) for the two CompCor variants: temporal (tCompCor) and anatomical (aCompCor). tCompCor components are then calculated from the top 2% variable voxels within the brain mask. For aCompCor, three probabilistic masks (CSF, WM and combined CSF+WM) are generated in anatomical space. The implementation differs from that of Behzadi et al. (2007) in that instead of eroding the masks by 2 pixels on BOLD space, the aCompCor masks are subtracted a mask of pixels that likely contain a volume fraction of GM. This mask is obtained by dilating a GM mask extracted from the FreeSurfer’s aseg segmentation, and it ensures components are not extracted from voxels containing a minimal fraction of GM. Finally, these masks are resampled into BOLD space and binarized by thresholding at 0.99 (as in the original implementation). Components are also calculated separately within the WM and CSF masks. For each CompCor decomposition, the k components with the largest singular values are retained, such that the retained components’ time series are sufficient to explain 50 percent of variance across the nuisance mask (CSF, WM, combined, or temporal). The remaining components are dropped from consideration. The head-motion estimates calculated in the correction step were also placed within the corresponding confounds file. The confound time series derived from head motion estimates and global signals were expanded with the inclusion of temporal derivatives and quadratic terms (Satterthwaite et al., 2013). Frames that exceeded a threshold of 0.5 mm FD or 1.5 standardised DVARS were annotated as motion outliers. The BOLD time-series were resampled into standard space, generating a preprocessed BOLD run in MNI152NLin2009cAsym space. First, a reference volume and its skull-stripped version were generated using a custom methodology of fMRIPrep. All resamplings can be performed with a single interpolation step by composing all the pertinent transformations (i.e. head-motion transform matrices, susceptibility distortion correction when available, and co-registrations to anatomical and output spaces). Gridded (volumetric) resamplings were performed using antsApplyTransforms (ANTs), configured with Lanczos interpolation to minimize the smoothing effects of other kernels (Lanczos, 1964). Non-gridded (surface) resamplings were performed using mri_vol2surf (FreeSurfer). Many internal operations of fMRIPrep use Nilearn (Abraham et al., 2014) 0.6.2 (RRID:SCR_001362), mostly within the functional processing workflow. The temporal signal-to-noise ratio was calculated for each voxels as the mean un-smoothed signal divided by the standard deviation across time.

### Regions of interest definition

For the hippocampus, participant-specific masks were obtained from FreeSurfer segmentations. For the entorhinal cortex, we used ASHS-derived segmentations of medial temporal lobe structures (Yushkevich et al., 2015), based on the U-Penn T1 atlas optimized for aging populations (Xie et al., 2019), which provides anatomically detailed delineation of medial temporal lobe cortices in older adults. Individual masks were transformed from native space to MNI space and resampled to the functional EPI resolution using FSL FLIRT. For posterior cingulate, we used a group-level mask of Brodmann Area 31 generated with the WFU PickAtlas toolbox (Maldjian et al., 2003) based on the Talairach atlas. This mask was resliced to the EPI dimensions using SPM Reslice.

To account for regional neurodegeneration, FreeSurfer-derived measures of atrophy in the hippocampus, entorhinal cortex, and posterior cingulate cortex were included as covariates in the multivariate analyses.

### fMRI analyses

#### Overview

All analyses were conducted using SPM12. Functional images were high-pass filtered with a cutoff of 180 seconds, to account for the slower nature of the self-pace task, with longer trials in some subjects. First-level models included nuisance regressors comprising the six realignment parameters (x, y, z, pitch, roll, yaw), their temporal derivatives, and quadratic terms. Additional regressors included motion outliers (defined as frames with framewise displacement exceeding 0.9), the mean signal extracted from either individual brain masks or anatomically defined, eroded CSF and white matter masks, non-steady-state volumes and the average temporal CompCor (tCompCor) as an estimate of physiological noise. Spatial smoothing was applied using a Gaussian kernel with an 8 mm full-width at half-maximum (FWHM). Voxels located outside the brain were excluded from all analyses.

#### Task-phase model

All trial phases were entered into the first-level models and the main effect of the retrieval phase, where participants are supposedly using memory to recall object locations and navigate to them, was subsequently evaluated.

#### Hexadirectional models

We used the Matlab toolbox GridCAT (Stangl et al., 2017) to estimate hexadirectional code metrics. Periods of active movement and their corresponding allocentric angles were smoothed over a one-second moving average window. Three models were estimated on two different partitions of the trajectory events, divided between the first (GLM1) and second half of events (GLM2-3), per run. In the first model (GLM1), trajectory event angles defined two regressors as follows: *sin*(*αt* ∗ 6) and *cos*(*αt* ∗ 6), where *αt* corresponds to the event angle. The mean axes orientation is calculated from the weighted average beta estimates (*β*1 and *β*2) across all voxels in subject-specific bilateral entorhinal cortex, as delineated by the ASHS segmentation. The orientation of the hexagonal axes was calculated for each run separately. Alignment to the axes orientation was then calculated with a parametric regressor (GLM2) by the following function *cos*[6(*α_t_* − *φ*)], with *α_t_* event angles and *φ* the mean axes orientation. Under these circonstances, a voxel beta estimate close to 1 indicates perfect alignment with the mean axes orientation (or its 60^◦^ multiples) and −1 indicates maximal misalignment (i.e., an offset of 30^◦^ to *φ*). Hexadirectional metrics could also be calculated based on two regressors (aligned or misaligned to the *φ*, averaged across 60^◦^ multiples, GLM3). GLM2 and GLM3 gave similar and coherent results, tough we report GLM2 only.

Individual first-level maps were the averaged in the entorhinal cortex ROI, or based on a second-level cluster identified in the *control group*. The local maxima was close to the entorhinal cortex (coordinate: [−22, 11, −42]). This coordinate was used to define a spherical mask (with 1.5 mm radius and 19 voxels, thus roughly corresponding to the group effect cluster size of 14 voxels), used to average the beta estimate for the rest of the participants.

Spatial and temporal stability of the hexadirectional pattern were calculated to assess the reliability of hexadirectional representations. Spatial stability was quantified as the consistency of voxel-wise axes orientations within the ROI using Rayleigh’s z statistic, with higher values indicating stronger clustering of orientations across voxels. Temporal stability was measured as the proportion of voxels whose axes orientation remained consistent across the first and second halves of the data (e.g., within ±15°), reflecting the stability of the hexadirectional pattern over time. Together, these metrics ensured that observed differences in hexadirectional activity were not due to spatial or temporal variability of the underlying representation.

### Object-centered model

Similar to Bécu et al. (2025), this model uses stick functions sampling each second of “active navigation” periods (defined by a velocity > 1vu/second). The parametric modulator for this regressor was the egocentric direction to the object in the range [0,180] degrees, rescaled onto [1,-1] range, with 1 representing moment in time where the object is directly in front of the participant, and −1 when the object is directly behind the participant. Subsequently, egocentric direction values were discretized into 8 regressors, as shown in figure 4E.

## Statistics

For each statistical model, raw data were transformed with a Box–Cox transformation to achieve normality (Osborne, 2010) and outlier data points (defined as lower/higher than 3 inter-quartile range of the 25*^th^* or 75*^th^* distribution percentile) were removed. Normality of the data is verified by visual inspection of Q–Q plots. When normality could not be attained, non-parametric alternatives, like the Wilcoxon rank test, were used.

For one-way ANOVA, the effect size was evaluated with the eta-square: 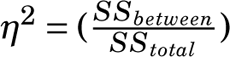 with *SS* the sum of square associated with the between and total effects. *η*^2^ indicates proportion of variance of the dependent variable explained by the independent variable. For two-way models with both between and within factors, the generalized eta-square (ges, *n^2^_g_*) is reported. *η*^2^ of 0.01, 0.06 and 0.14 are considered small, medium and large effects, respectively. Alpha level for statistical significance was set at *p* < 0.05. Posthoc tests were evaluated with a Bonferroni correction. Exact p-values are reported, until *p* < 0.0001. Statistical tests were one-sided.

For the multivariate analyses, we conducted Bayesian multiple linear regression analyses using the brms package (version 2.23) in R, which uses Markov Chain Monte Carlo (MCMC) sampling. The dependent variable was the average beta estimate in each of our ROIs. Predictors included biomarker variables (amyloid, ptau, *APOE* status), behavioral measures (CERAD zscore, drop error), and covariates (age, sex, atrophy, tSNR). Multi-collinearity among predictors was assessed using variance inflation factors (VIF) derived from an equivalent linear model. Because of a strong and expected correlation between ptau and ttau, we decided to remove ttau from the modelling. All other variables had values below 3 indicating acceptable levels of collinearity. Continuous variables were standardized prior to analysis to facilitate interpretation of regression coefficients. Weakly regularizing priors were specified for regression coefficients (normal(0, 0.4)), and a Student-t prior was used for the residual standard deviation. Models were estimated using four MCMC chains with 8,000 iterations per chain (4,000 warm-up), and convergence was assessed using the potential scale reduction statistic (*R^* > 1.01), effective sample size (ESS), and inspection of trace plots. No convergence issues were observed. Posterior means and 95% credible intervals (CrI) are reported. Directional hypotheses were evaluated using posterior probabilities. Effects with posterior probability > 0.975 were interpreted as strong evidence for the hypothesized direction, probabilities between 0.90 and 0.975 were considered moderate evidence, and probabilities < 0.90 were considered weak or inconclusive evidence.

## Acknowledgments

The authors wish to thank Lene Pålhaugen and the DDI study teams at Akershus University Hospital and St. Olavs Hospital for their assistance with patient recruitment and examination.

## Author contributions

C.F.D., T.N.S, G.R.G, conceived the study. C.F.D., T.N.S, G.R.G, T.B., J.A.J., P.S., A.B., T.F, conceptualized the study. J.A.J, G.R.G, S.B.S recruited participants and acquired the data. M.B., T.B., J.A.J., G.R.G. curated the data. M.B., I.M.K, J.A.J. analyzed the data. M.B., T.B, J.A.J. and C.F.D. wrote and revised the manuscript. T.B, T.F., C.F.D supervised the project.

## Data and materials availability

The fMRI data underlying the results of this study were collected in compliance with Norwegian regulations governing data protection and privacy. Because of legal and ethical restrictions set by the Norwegian Data Protection Authority (Datatilsynet) and the Regional Committees for Medical and Health Research Ethics (REK), the raw imaging data cannot be made publicly available. Access to these data is restricted to approved research collaborators who have obtained the required ethical permissions and data-processing agreements in Norway. Researchers interested in accessing the data may contact the corresponding author to discuss possible collaborations within these limitations. Brain data (i.e. group-level brain maps, individual beta values) and behavioral data (i.e. task performance measures) supporting figures 1-4 will be deposited in the following repository at the time of publication: https://osf.io/ac4w9/. The group (CTR, PRE, EAD) will be shared on the repository but individual biomarker concentration will not be made publicly available. The MATLAB code used to model and analyze the fMRI data, along with the scripts used for behavioral analyses, will be made available alongside the data.

## Supplementary Material

## Supplementary Figures

**Figure S1.**
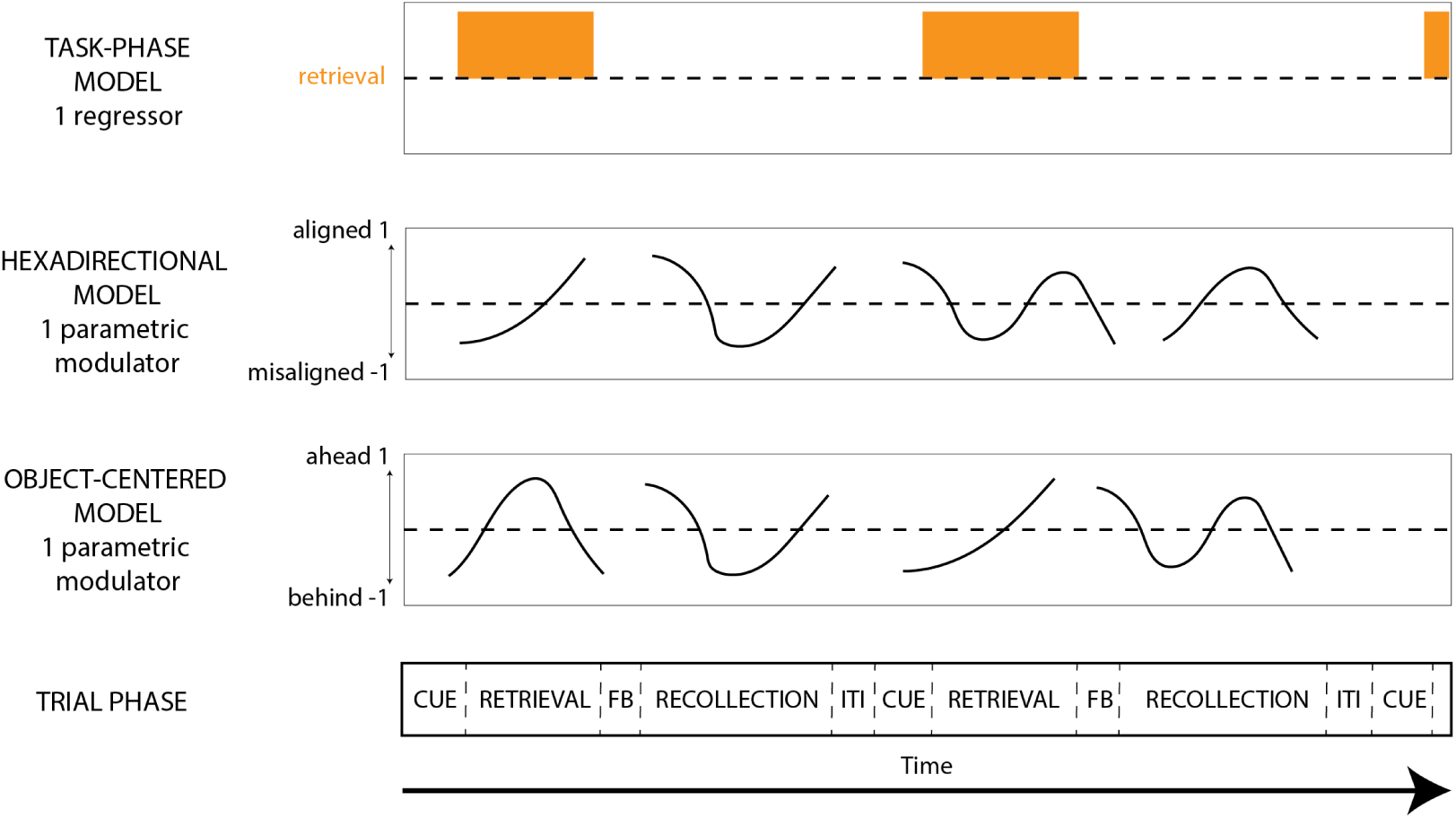
fMRI data modeling scheme. (Top) The task-phase model evaluated the main effect of the retrieval phase. Other trial phases were included in the model but were not analyzed further. (Middle) The hexadirectional model evaluated whether brain activity increased when participant trajectories were aligned with the putative axes orientation. A single parametric modulator was used to model trajectory direction. Stationary periods were not modeled. (Bottom) The object-centered model used a single parametric modulator to assess whether brain activity increased when the object was directly in front of the participant. ITI = inter-trial interval. FB = feedback

**Figure S2.**
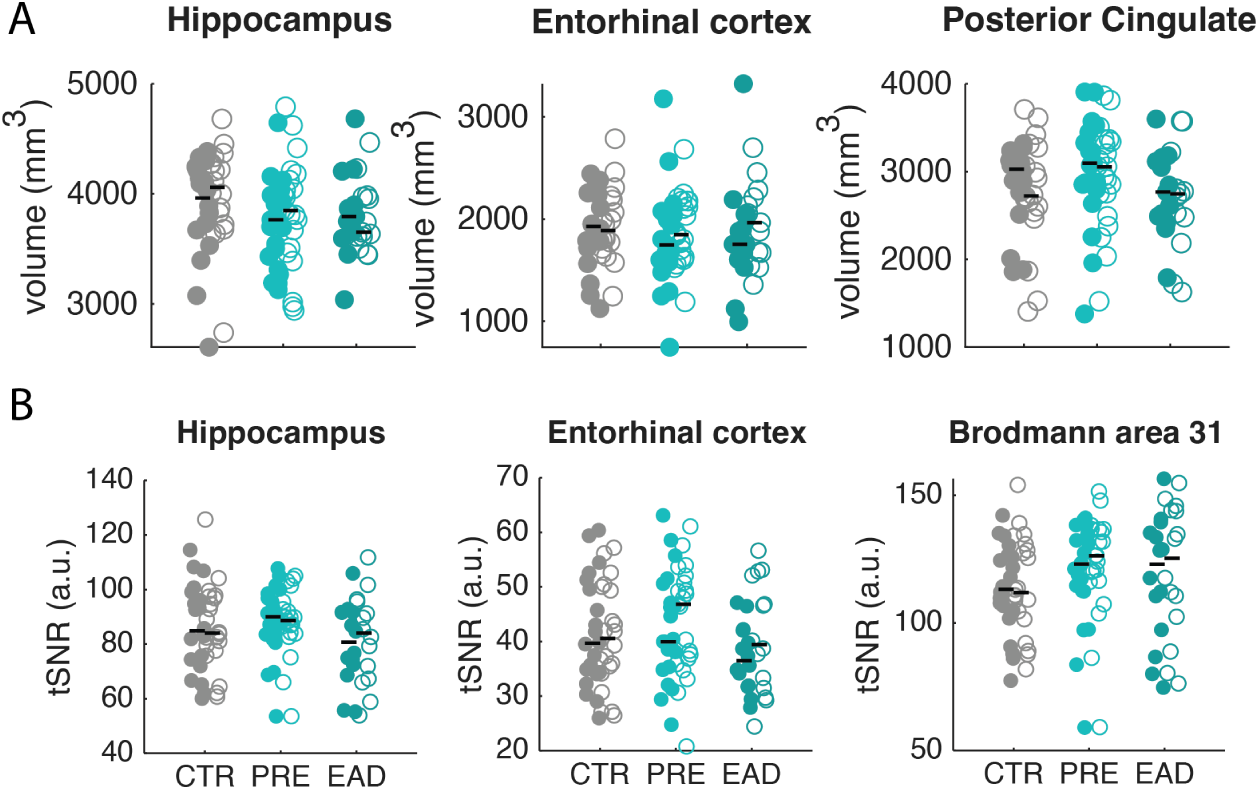
Atrophy and signal quality in the regions of interest. (A) Atrophy, measured as the grey matter volume within areas of interest, segmented by FreeSurfer. (B) Signal quality measured as temporal signal-to-noise ratio in individual masks (a.u., arbitrary unit).

**Figure S3.**
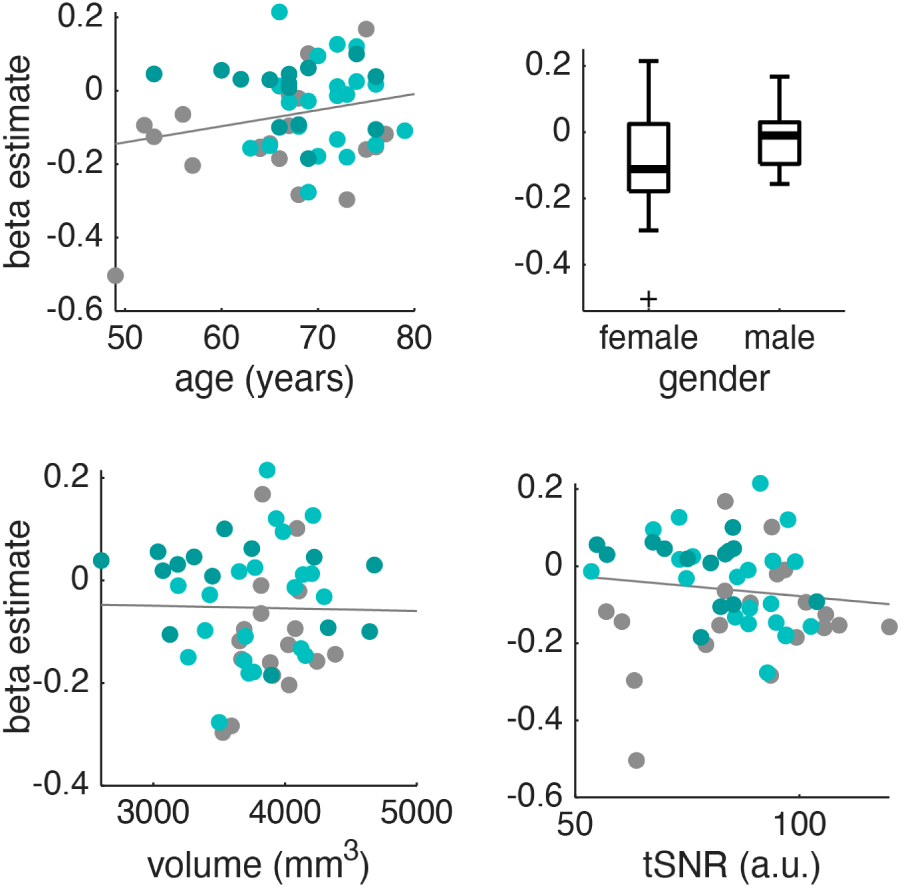
Association between hippocampal hyperactivity and the model covariates. **a.u.** = arbitrary unit

**Figure S4.**
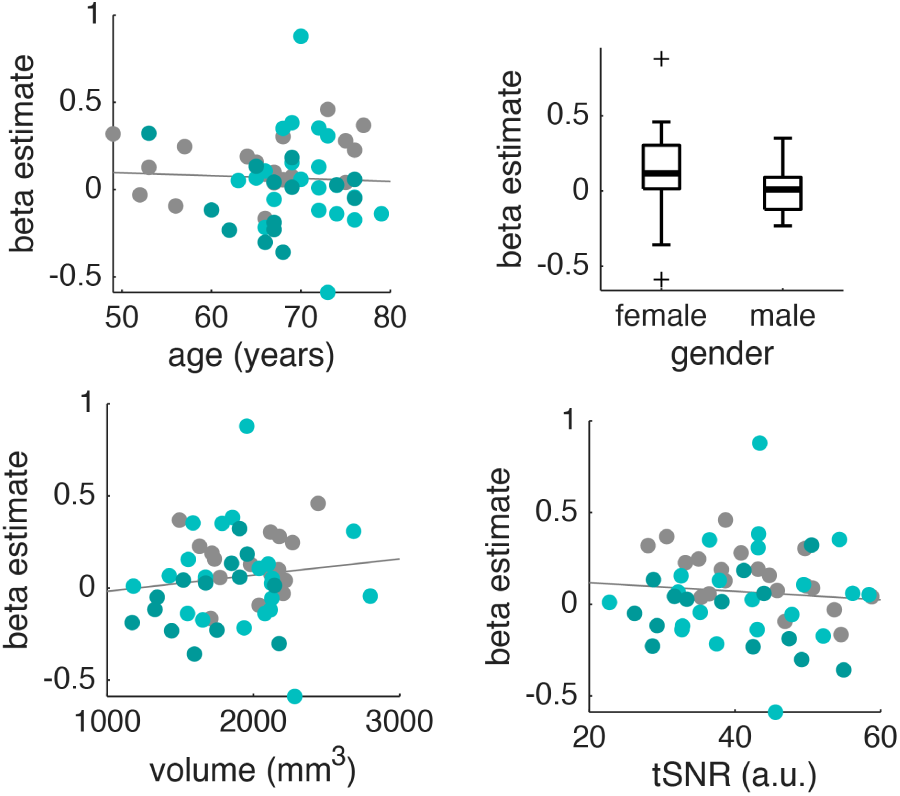
Association between hexadirectional activity and the model covariates. **a.u.** = arbitrary unit

**Figure S5.**
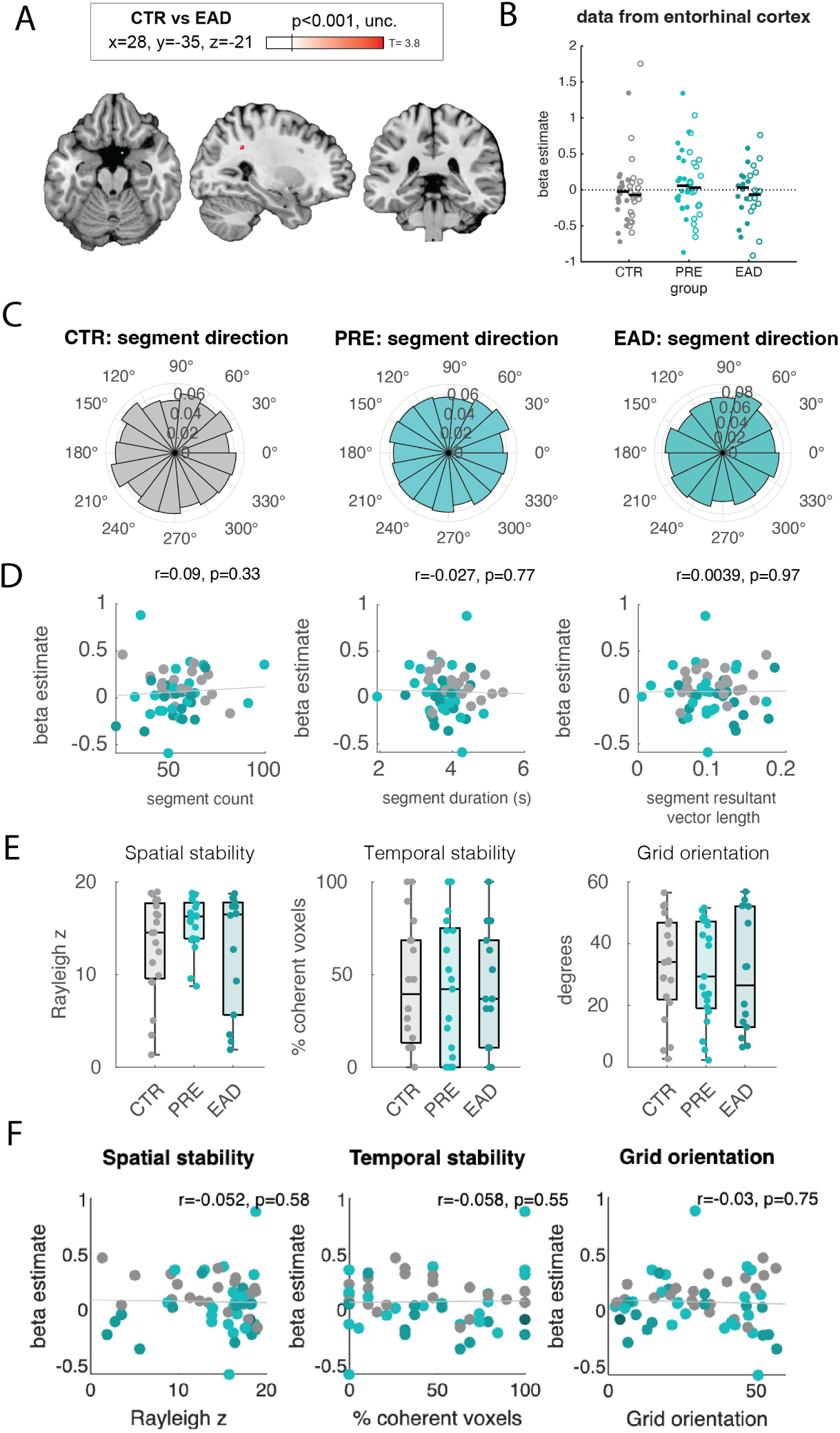
Control analyses of the hexadirectional modulation. (A) Whole-brain ANOVA comparing CTR to EAD at uncorrected threshold (unc.). Maximum T-score, and MNI slice location. (B) Beta estimate averaged in the entorhinal cortex. (C) Density-normalized histogram of trajectory segment direction, in allocentric space, for the three groups. CTR = controls, PRE = preclinical group, EAD = early AD group. (D) Correlations between the hexadirectional effect, averaged in the sphere of interest and segment count, duration (in seconds) and resultant vector length of segment directions. (E) Spatial, temporal stability and orientation of the putative axes between groups. (F) Correlations between the hexadirectional effect, averaged in the sphere of interest and spatial, temporal stability and orientation of the putative axes.

**Figure S6.**
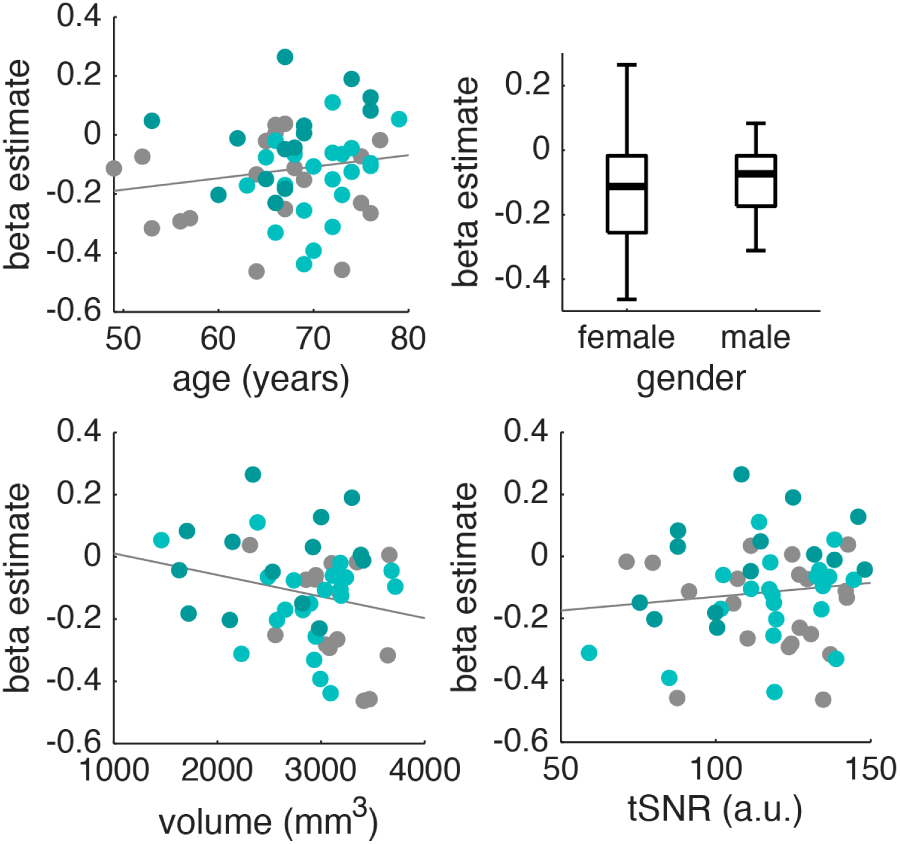
Association between object representations and the model covariates. **a.u.** = arbitrary unit

## Supplementary Tables

**Table S1.** Posterior estimates from Bayesian regression predicting hippocampal hyperactivity (VD). Estimates represent standardized regression coefficients. CrI = 95% credible interval; Post. Prob = posterior probability for the hypothesized direction. ** = strong evidence; * = moderate evidence.

| Predictor | Hypothesis | Estimate | Error | CrI Lower | CrI Upper | Post. Prob | Evidence |
| --- | --- | --- | --- | --- | --- | --- | --- |
| abeta_ratio | < 0 | -0.53 | 0.26 | -0.96 | -0.09 | 0.98 | ** |
| ptau | < 0 | -0.08 | 0.14 | -0.32 | 0.16 | 0.72 |  |
| APOE | < 0 | 0.05 | 0.23 | -0.33 | 0.43 | 0.41 |  |
| CERAD_zscore | < 0 | -0.05 | 0.16 | -0.31 | 0.21 | 0.61 |  |
| drop_error | < 0 | 0.16 | 0.15 | -0.08 | 0.41 | 0.13 |  |
| age | < 0 | -0.16 | 0.14 | -0.38 | 0.07 | 0.88 |  |
| sex | > 0 | 0.33 | 0.23 | -0.06 | 0.71 | 0.92 | * |
| tsnr | > 0 | -0.04 | 0.13 | -0.25 | 0.17 | 0.39 |  |
| atrophy | < 0 | 0.09 | 0.13 | -0.13 | 0.31 | 0.25 |  |

**Table S2.**
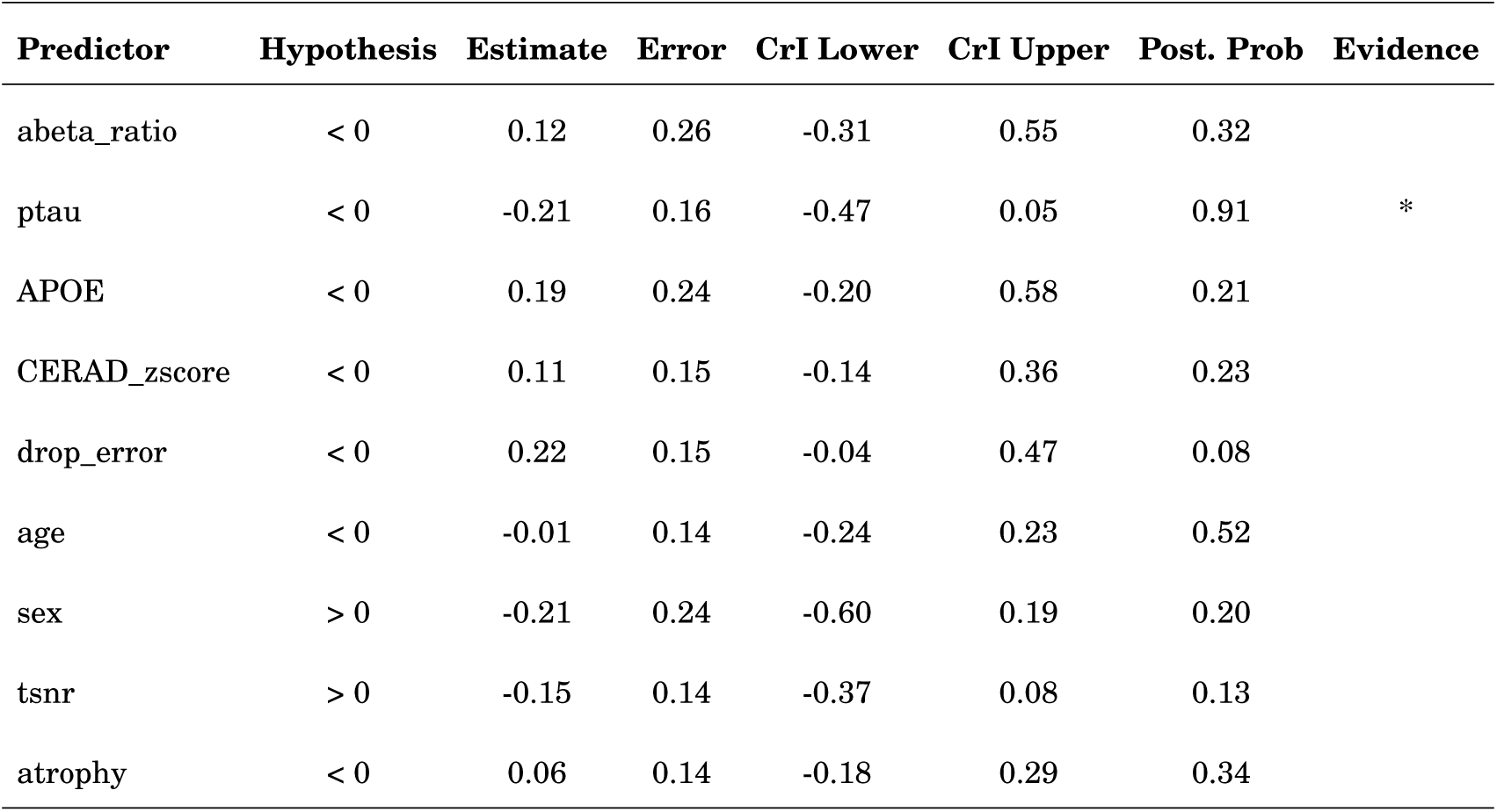
Posterior estimates from Bayesian regression predicting the hexadirectional modulation (VD). Estimates represent standardized regression coefficients. CrI = 95% credible interval; Post. Prob = posterior probability for the hypothesized direction. ** = strong evidence; * = moderate evidence.

| Predictor | Hypothesis | Estimate | Error | CrI Lower | CrI Upper | Post. Prob | Evidence |
| --- | --- | --- | --- | --- | --- | --- | --- |
| abeta_ratio | < 0 | 0.12 | 0.26 | -0.31 | 0.55 | 0.32 |  |
| ptau | < 0 | -0.21 | 0.16 | -0.47 | 0.05 | 0.91 | * |
| APOE | < 0 | 0.19 | 0.24 | -0.20 | 0.58 | 0.21 |  |
| CERAD_zscore | < 0 | 0.11 | 0.15 | -0.14 | 0.36 | 0.23 |  |
| drop_error | < 0 | 0.22 | 0.15 | -0.04 | 0.47 | 0.08 |  |
| age | < 0 | -0.01 | 0.14 | -0.24 | 0.23 | 0.52 |  |
| sex | > 0 | -0.21 | 0.24 | -0.60 | 0.19 | 0.20 |  |
| tsnr | > 0 | -0.15 | 0.14 | -0.37 | 0.08 | 0.13 |  |
| atrophy | < 0 | 0.06 | 0.14 | -0.18 | 0.29 | 0.34 |  |

**Table S3.** Posterior estimates from Bayesian regression predicting the object directional activity (VD). Estimates represent standardized regression coefficients. CrI = 95% credible interval; Post. Prob = posterior probability for the hypothesized direction. ** = strong evidence; * = moderate evidence.

| Predictor | Hypothesis | Estimate | Error | CrI Lower | CrI Upper | Post. Prob | Evidence |
| --- | --- | --- | --- | --- | --- | --- | --- |
| abeta_ratio | < 0 | -0.20 | 0.27 | -0.65 | 0.25 | 0.77 |  |
| ptau | < 0 | -0.12 | 0.15 | -0.37 | 0.12 | 0.80 |  |
| APOE | < 0 | -0.25 | 0.24 | -0.64 | 0.14 | 0.86 |  |
| CERAD_zscore | < 0 | -0.45 | 0.15 | -0.69 | -0.20 | 1.00 | ** |
| drop_error | < 0 | -0.05 | 0.16 | -0.31 | 0.22 | 0.62 |  |
| age | < 0 | 0.11 | 0.14 | -0.12 | 0.34 | 0.22 |  |
| sex | > 0 | -0.03 | 0.25 | -0.44 | 0.38 | 0.46 |  |
| tsnr | > 0 | 0.25 | 0.14 | 0.02 | 0.48 | 0.96 | * |
| atrophy | < 0 | -0.18 | 0.13 | -0.40 | 0.04 | 0.91 | * |

